# An SR45-dependent exonic splicing enhancer is essential for regulating alternative splicing in Arabidopsis

**DOI:** 10.1101/2025.09.27.678957

**Authors:** Yawen Zhu, WenYujie Shi, Liqun Zhang, Fangsheng Liao, Keyi Yang, Jiaqi Wang, Hui Zhang, Jirong Huang, Weihua Huang

## Abstract

Alternative splicing (AS) is a critical gene regulatory mechanism that underpins developmental plasticity and environmental adaptation in eukaryotes. Although numerous *cis*-elements and *trans*-factors governing AS have been characterized in animals, their functional roles in plants remain largely unexplored. We previously identified the *Arabidopsis sot5* mutant, which exhibits leaf virescence due to a mutation in the conserved 5’ splice site (5’ss) of intron 7. Here, through characterization of a suppressor of *sot5* (*E61*), we found that *E61* harbors a G to A mutation in exon 8 of the *SOT5* gene. Interestingly, this mutation creates a synonymous codon but disrupts the exonic splicing enhancer (ESE) motif (AAGAAGA), thereby reducing the splicing efficiency of the adjacent intron 7. Retention of this intron produces a functional protein that rescues the *sot5* virescent phenotype. We confirmed the causal role of this ESE by using a cytidine base editor (CBE) to convert the ESE sequence to AAGAAAA or AAAAAAA in *sot5*. The edited plants phenocopied the suppressor *E61*. Genetic analyses revealed that mutations of *SR45* encoding an RNA-binding protein but not its paralog *SR45a* similarly rescues the *sot5* phenotype by impairing intron 7 splicing. Furthermore, we demonstrated that SR45 directly binds to the ESE motif *in vitro*, and that mutating the proximal ESEs in intron 2 of another SR45 target, *GPDHC1* (*Glycerol-3-Phosphate Dehydrogenase C1*), recapitulates the splicing defect observed in the *sr45* mutant. These findings establish that SR45 specifically recognizes AAG-repeat ESEs to enhance the splicing of adjacent weak introns, revealing a potent strategy for fine-tuning gene expression through targeted editing of splicing regulatory elements for crop improvement.

## Introduction

Alternative splicing (AS) is a prevalent gene expression regulatory mechanism that enables a single gene to yield multiple distinct RNA products in eukaryotes (Lee and Rio, 2015). RNA splicing is carried out by the spliceosome, a large complex consisting of the pre-mRNA substrate and over 180 proteins, which recognizes canonical *cis*-regulatory elements including the conserved 5′ and 3′ splice sites (5′ss and 3′ss), the branch point (BP), and the polypyrimidine tract (Shao et al., 2012; Wilkinson et al., 2020; Zhu et al., 2020). The efficiency of splice site recognition is a key determinant for AS outcomes. It is well documented that strong consensus sites facilitate constitutive splicing, while weak splice sites are selectively used and highly influenced by auxiliary *cis*-elements located within adjacent exonic or intronic regions (Fairbrother et al., 2002; Wang et al., 2004; Chen and Manley, 2009). These auxiliary elements, known as exonic/intronic splicing enhancers (ESEs/ISEs) or silencers (ESSs/ISSs), are bound by *trans*-acting splicing factors, such as serine/arginine-rich (SR) proteins (Fu, 1993; Yeakley et al., 1996; Bourgeois et al., 2004) and heterogeneous nuclear ribonucleoproteins (hnRNPs, Rothrock et al., 2005; Geuens et al., 2016), which promote and repress splice site selection, respectively. The expression of splicing factors varies across different cell types and environmental conditions, allowing for precise spatiotemporal control of AS.

SR proteins conserved in all plants and metazoans are characterized by one or two RNA recognition motifs (RRMs) at the N-terminal and an arginine/serine dipeptide-rich (RS) domain mediating protein–protein interaction at C-terminal. They have been shown to play roles in constitutive and alternative splicing, mRNA export and translation. For example, during early spliceosome assembly, SR proteins stabilize U1 small nuclear ribonucleoprotein (snRNP) binding at the 5′ss through interactions with U1-70kDa protein (U1-70K). They also assist U2 small nuclear ribonucleoprotein auxiliary factor 35 kDa subunit (U2AF35) in facilitating the recognition and binding of U2 snRNP to the 3′ss (Bourgeois et al., 2004; Chen and Manley, 2009). Remarkably, another group of proteins, called SR-like proteins usually containing a single RRM domain flanked by two RS domains, resemble the SR protein in structure and functions related to RNA processing. Among the best studied of the SR-like splicing factors is the transformer 2 (Tra2) protein, which regulates sex determination by binding splicing enhancer of pre-mRNA from the sex determination gene *doublesex* (*dsx*) in *Drosophila* (Hedley and Maniatis, 1991; Tian and Maniatis, 1993). Its human homolog Tra2β binds to the ESE sequence (AAAGAAGGAAGG) in exon 7 of the *survival motor neuron* (*SMN2*) gene. This binding promotes the inclusion of exon 7, stimulating the production of full-length functional protein. This process compensates for the loss of *SMN1* function and can alleviate symptoms in patients with spinal muscular atrophy (SMA) (Tacke et al., 1998; Hofmann et al., 2000).

Compared to those in animals, the roles of SR- and SR-like proteins and their target enhancers in alternative splicing remain limited in plants. Arabidopsis has eighteen canonical SR proteins and two SR-like proteins (Barta et al., 2010). Many of these proteins share the same RNA-binding motifs with their animal counterparts and play similar roles in enhancing splicing (Lazar et al., 1995). Arabidopsis SR45, a homolog of human RNPS1, and SR45a, a homolog of Tra2β, are critical regulators in growth, development and stress responses (Ali et al., 2007; Xing et al., 2015; Zhang et al., 2014; Carvalho et al., 2016; Zhang et al., 2017; Albaqami et al., 2019; Fanara et al., 2022; Albaqami, 2023; Albuquerque-Martins et al., 2023; Albaqami et al., 2019; Li et al., 2021). Recently, AtSR45 was reported to be the effector of Glomeromycotina-specific SP7 that is involved in symbiosis with arbuscular mycorrhizal (AM) fungi (Betz et al., 2024). AtSR45 specifically binds the core RNA motif AAGAAG in both intron-containing and intronless genes (Xing et al., 2015; Fanara et al., 2024), and directly interacts with the core splicing components U1-70K and U2AF35 (Day et al., 2012). Despite its known RNA-binding specificity and physiological importance, direct evidence linking AtSR45 to a specific cis-regulatory element that controls a defined splicing event has been lacking.

Our previous work established a powerful genetic system to study splice site selection by screening suppressors for *sot5* (*suppressor of thylakoid formation 5*) in Arabidopsis. The *sot5* mutant, which harbors a point mutation at the 5′ss of intron 7 in a pentatricopeptide repeat (PPR) gene, exhibits a virescent leaf phenotype. The point mutation leads to activation of two proximal cryptic splice sites and retention of intron 7, generating three additional mRNA variants (Huang et al, 2018). We demonstrated that mutations in two U1 snRNP components, RBP45d and PRP39a, rescue the virescent leaf phenotype of *sot5* via upregulating the expression of a cryptically spliced variant that encodes a mutated but functional SOT5 protein (Huang et al, 2022), revealing that RBP45d and PRP39a contribute to the accurate selection of specific 5′ss during spliceosome assembly. Thus, the readily observable leaf-color phenotype makes the *sot5* suppressor system a powerful tool for dissecting the regulatory mechanisms of alternative splicing.

Here, through the characterization of an intragenic suppressor (*E61*) of the *sot5* mutant, we identify an AAGAAG motif in exon 8 of *SOT5* that functions as an ESE to promote splicing of the upstream weak intron. We demonstrate that loss-of-function mutation in *SR45* phenocopies the *E61* suppressor and rescues the *sot5* phenotype. Furthermore, we confirm that SR45 protein binds directly to this specific ESE motif *in vitro*. Our findings provide direct evidence that SR45 regulates AS by binding a specific splicing enhancer element, elucidating a key mechanism of splicing regulation in plants.

## Results

### An intragenic mutation suppresses the virescent phenotype of *sot5* by reducing the splicing efficiency of intron 7

In a screen for suppressors of the *sot5* mutant using ethyl methanesulfonate (EMS)-mutagenized seeds, we identified the *E61* suppressor line exhibiting a wild-type (WT) phenotype (Figure 1A). The chlorophyll content and maximum quantum efficiency of photosystem II (PSII) *F*_v_/*F*_m_ values of *E61* were comparable to WT and significantly higher than those of *sot5* (Figure 1B and 1C). To determine the molecular mechanism underlying the suppression, we analyzed splicing patterns of the mutated *SOT5* gene in *E61*. RT-PCR analysis revealed a marked increase in transcripts retaining intron 7 and a corresponding decrease in transcripts utilizing the -22 nt cryptic splice site in *E61*, compared to *sot5* (Figure 1D). Indeed, quantitative PCR results confirmed a significant reduction in the splicing efficiency of intron 7 in *E61* (Figure 1E). These results indicate that suppression of the *sot5* phenotype in *E61* is attributed to intron 7 retention.

**Figure 1.**
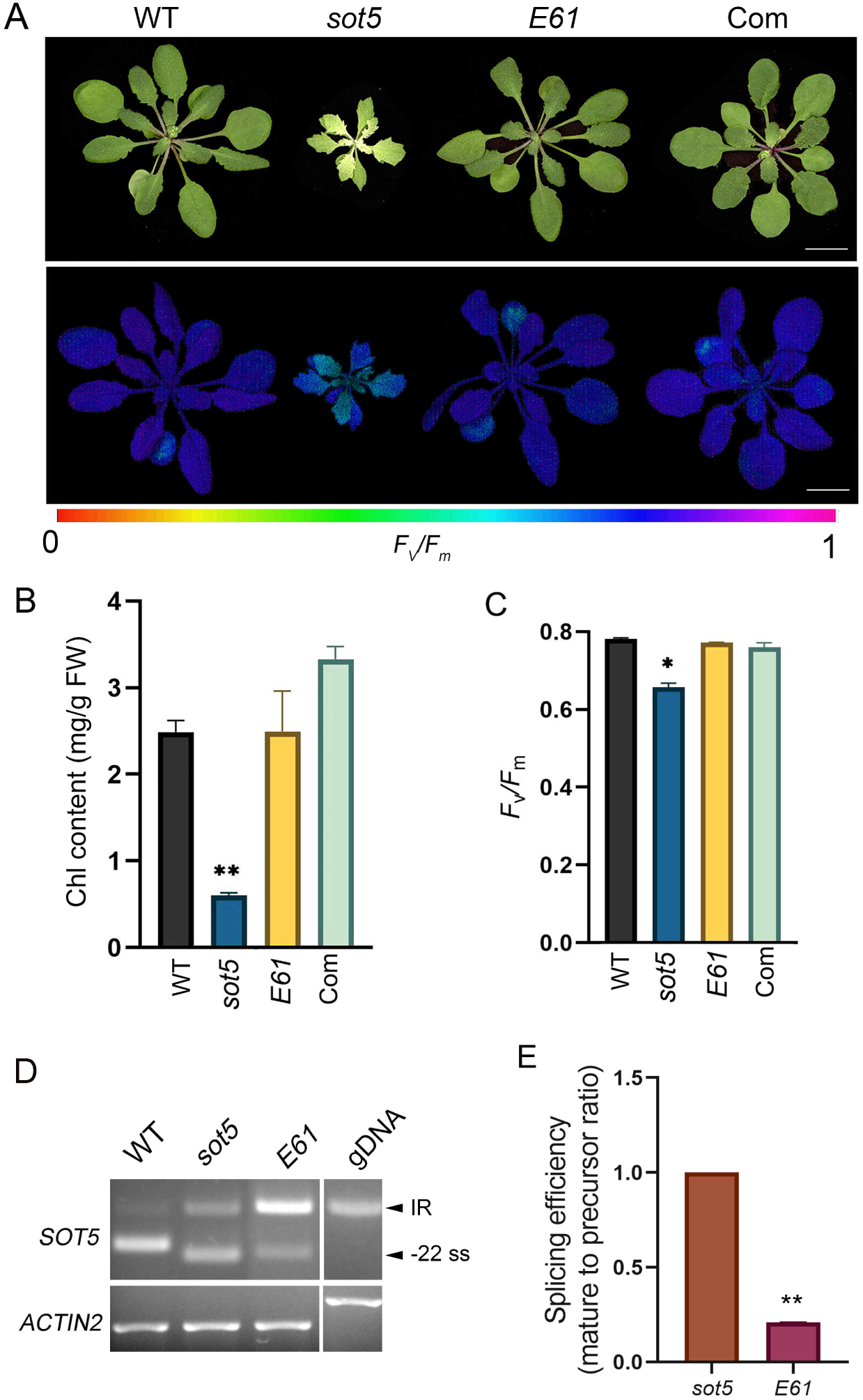
Characterization of a suppressor of *sot5*, *E61*. (A) Phenotype and Chlorophyll (Chl) fluorescence image of *F*_v_/*F*_m_ of 35-day-old WT, *sot5, E61* and *E61* complementation plants. Bar = 1 cm. (B) Chl content of the leaves from plants shown in (A). (C) Quantification of *F*_v_/*F*_m_ in (A). The data represent means ± SD of three biological replicates. Asterisks indicate significant differences between WT and *sot5* (Student’s *t* test). *, *P* < 0.05; **, *P* < 0.01. (D) RT-PCR analysis of the splicing variants of *SOT5* intron 7 in *sot5* and *E61*. IR indicates the transcript retained intron 7. -22 ss indicates the transcript derived from -22 cryptic splice site. The RT-PCR primers were indicated by the red arrows in Figure 2A. (E) Quantification of *SOT5* intron 7 splicing efficiency (spliced transcripts/unspliced transcripts) by qPCR. The RT-qPCR primers were indicated by the green arrows in Figure 2A. The splicing efficiency in the *sot5* mutant was set as 1. The values are means of three biological replicates (bars indicate SD). Asterisks indicate significant differences between WT and *sot5* (Student’s *t* test). **, *P* < 0.01.

To clone the *E61* gene, we first investigated whether the phenotypic rescue was due to a second-site mutation within the *SOT5* gene. Sanger sequencing identified a G-to-A point mutation at the 9th nucleotide of exon 8, proximal to the 3′ss of intron 7, indicating that *E61* is an intragenic suppressor of *sot5* (Figure 2A). Genetic analysis of a cross between *sot5* and *E61* showed that F_1_ plants displayed an intermediate phenotype (Figure S1), and the F_2_ generation segregated into WT-like, intermediate, and *sot5*-like phenotypes in an approximate 1:2:1 ratio (Table S1). These results suggest that *E61* acts in a semi-dominant manner in the *sot5* background. Additionally, no *sot5*-like plants were observed in the F_2_ population from a cross between *E61* and Col-0, confirming that suppression is caused by the identified intragenic mutation.

**Figure 2.**
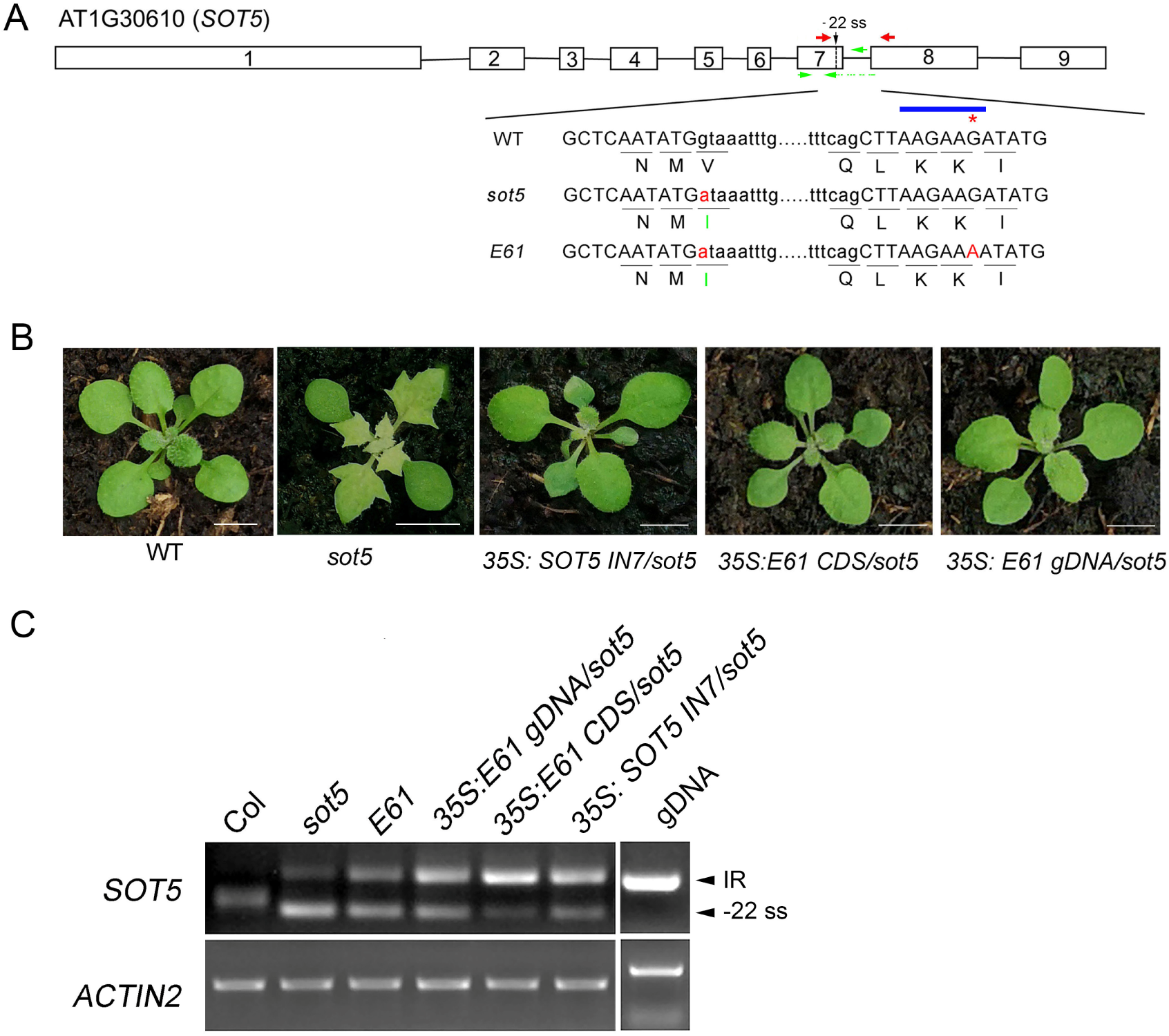
*E61* harbors a synonymous exonic mutation reducing intron 7 splicing. (A) Schematic diagram of *SOT5* gene structure with mutation sites in *sot5* and *E61*. The boxes indicate exons and the black lines indicate introns. Black dashed line indicates the cryptic splice site, -22 ss, that was activated in *sot5* mutant. Red and green arrows indicate primers used for RT-PCR and RT-qPCR analysis, respectively. Uppercase letters represent exon sequences, and lowercase letters represent intron sequences. The blue line marks the putative exonic splicing enhancer (ESE) motif. The red asterisk denotes the G-to-A mutation within ESE in *E61*. Translated amino acids are shown below; green letters indicate mutated residues. (B) Complementation assays of *sot5*. Representative transgenic plants overexpressing *E61* gDNA, *E61* CDS (*E61* retaining intron 7), and *sot5 IN7* CDS (*SOT5* retaining intron 7) in the *sot5* background. Bar = 1 cm. (C) RT-PCR analysis of *SOT5* intron 7 splicing variants from plants shown in (B).

To validate the intragenic mutation, we transformed the *sot5* mutant with the *E61* CDS (retaining intron 7) or *E61* genomic DNA (gDNA) driven by the 35S promoter. Both constructs fully complemented the *sot5* phenotype (Figure 1A, 2A), suggesting that the product, a 1006-amino acid polypeptide, encoded by the *E61* gene is functional. Notably, the G-to-A mutation in exon 8 is synonymous and does not alter the amino acid sequence (Figure 2A). We then tested whether the intron 7-retaining *SOT5* cDNA (designated *SOT5 IN7* CDS) itself could produce a functional protein. Our results showed that overexpression of *SOT5 IN7* CDS fully complemented in the *sot5* phenotype (Figure 2B). RT-PCR confirmed overexpression of the respective transgenes (Figure 2C). These results confirmed our previous presumption that the intron 7-retaining *SOT5* transcript encodes a mutated but functional SOT5 containing 10 PPR motifs (Huang et al., 2018). Taken together, the above results demonstrate that the *E61* suppressor harbors a synonymous point mutation in exon 8 of *SOT5*. This mutation reduces the splicing efficiency of intron 7, leading to the accumulation of a new functional SOT5 protein isoform retaining 10 PPR motif.

### The intragenic mutation disrupts an ESE in exon 8 of *SOT5* and the ESE functions only when a weak splice site is present

ESEs are often purine-rich motifs (A/G) located in exons near the 5′ss or 3′ss, and serve as binding platforms for SR and SR-like proteins. We found a candidate ESE motif within exon 8 of *SOT5*, spanning nucleotides from 4 to 10 (AAGAAGA) (Figure 2A). The *E61* mutation lies directly within this element (at the 9th nucleotide of exon 8), suggesting that the mutation disrupts ESE function, thereby reducing the splicing efficiency of the upstream intron 7. To verify this hypothesis, we used a Cytosine Base Editor (CBE) to introduce G-to-A mutations at position 6 and 9 of exon 8 within this putative ESE in *sot5* backgrounds. Notably, these mutations do not alter the protein coding sequence. We obtained two independent homozygous edited lines, *sot5 EX8-CR-1* and *sot5 EX8-CR-2*, which exhibited the WT-like phenotype (Figure 3A, 3B and 3C). As a positive control, we also introduced a G-to-A mutation at the 3′ss of *SOT5* intron 7 in the *sot5* mutant using CBE. Our results showed that the resulting *sot5 IN7-CR* plants also displayed the WT-like phenotype (Figure 3A, 3B and 3C). RT-PCR analysis revealed that simultaneous disruptions of the 5′ss and 3′ss of intron 7 in *sot5 IN7-CR* plants severely impaired the splicing of intron 7 (Figure 4A). Interestingly, the splicing efficiency of intron 7 was similarly reduced in *sot5 EX8-CR-1* and *sot5 EX8-CR-2* lines (Figure 4C), demonstrating that ESE mutations within exon 8 mimic the effect of direct mutation in a core splice site.

**Figure 3.**
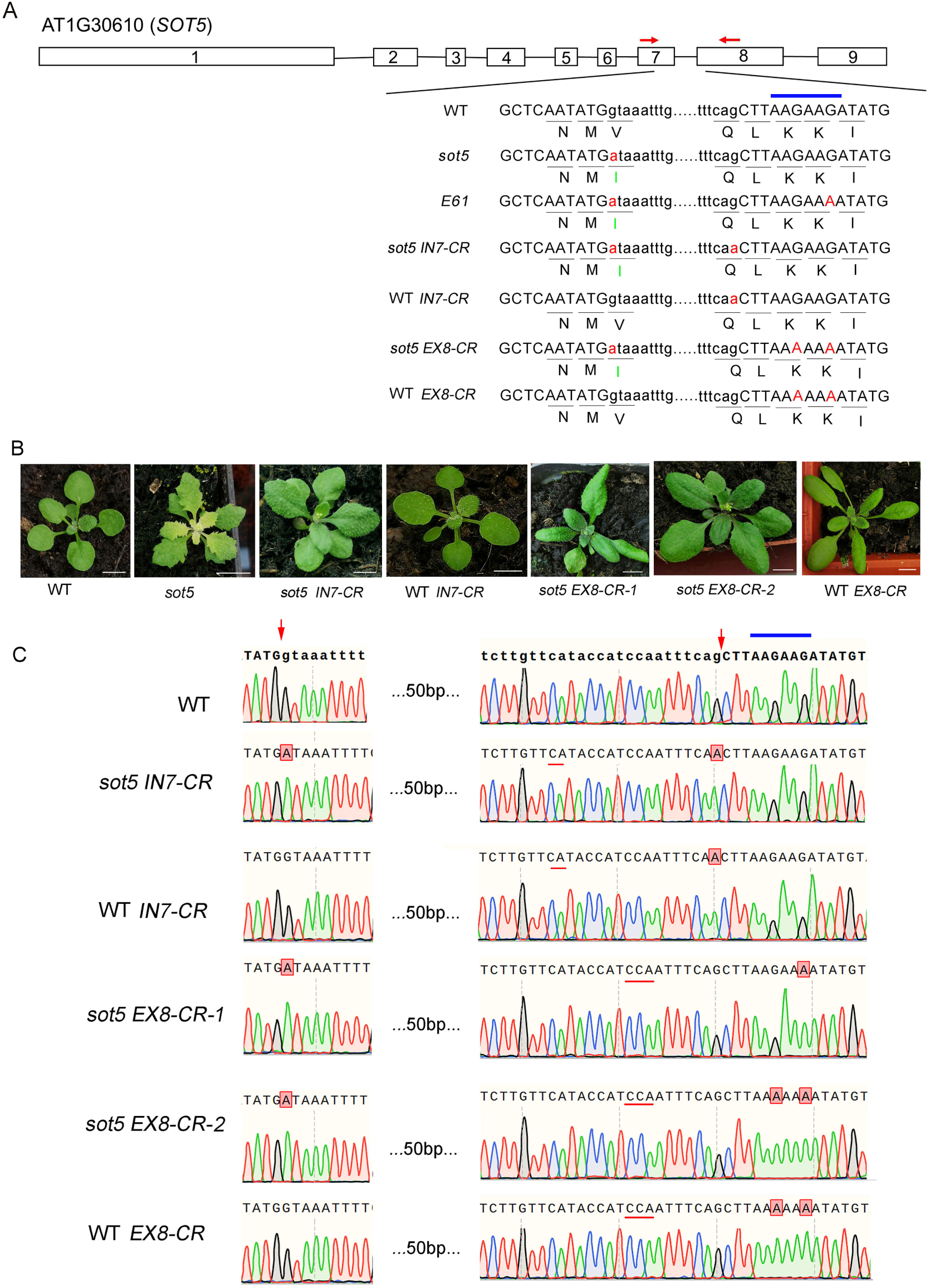
Disruption of an ESE in *SOT5* exon 8 recapitulates the *E61* suppressor phenotype. (A) Schematic diagram of the *SOT5* gene structure with mutation sites indicated. Red for original *sot5* and *E61* mutations, and the targeted sites in CBE-edited plants, *sot5-IN7-CR,* WT*-IN7-CR, sot5-EX8-CR* and WT*-EX8-CR*. The blue line marks the ESE motif. (B) Phenotype of the CBE-edited, WT and *sot5* plants. Bar = 1 cm. (C) Sanger sequencing chromatograms around the targeted sites. The red lines indicate the protospacer adjacent motif (PAM) sequence. The red arrows indicate the 5’ and 3’ splicing border of *SOT5* intron 7. The blue line marks the ESE motif.

**Figure 4.**
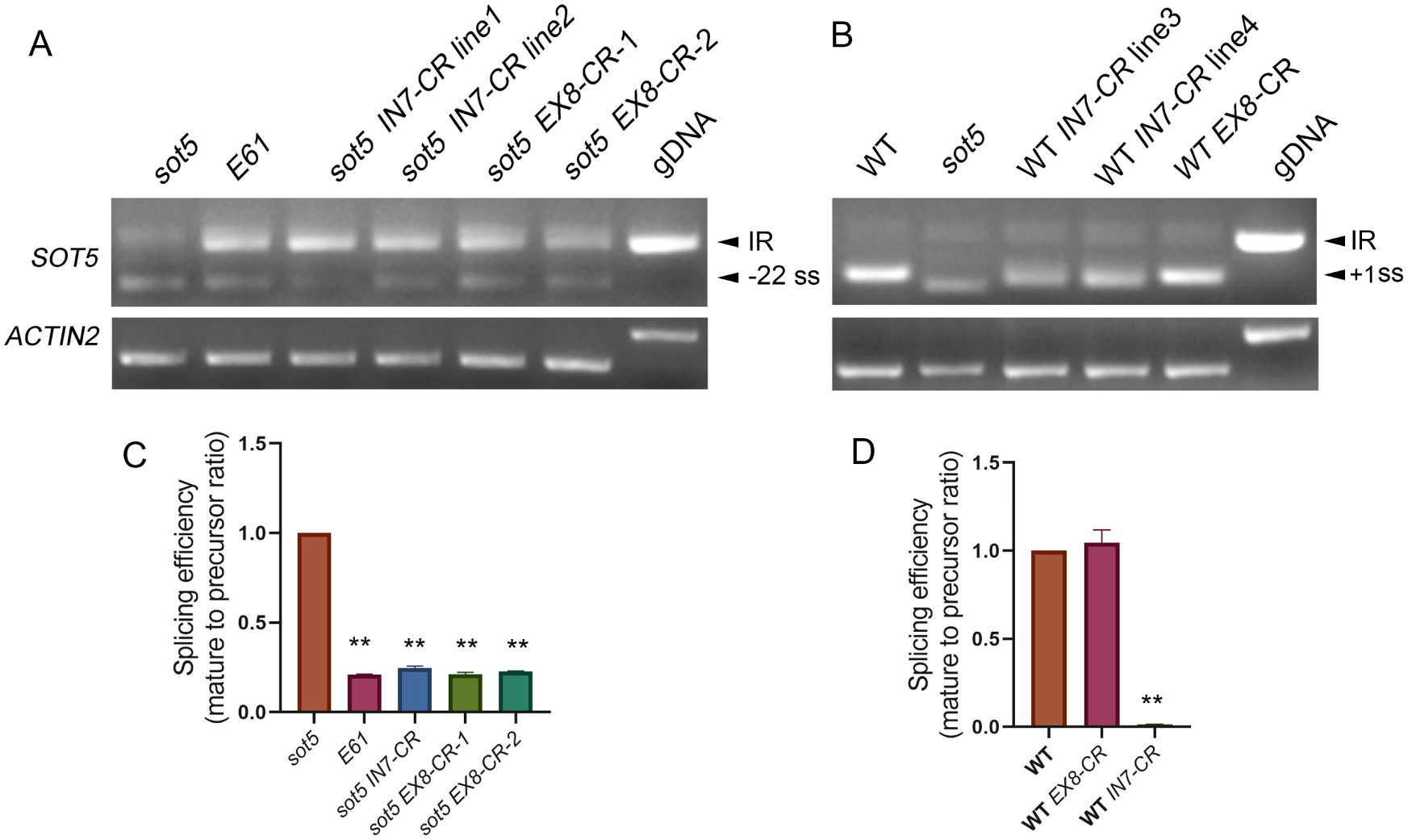
Splicing efficiency analysis of *SOT5* intron 7. Semi-quantitative PCR analysis of *SOT5* intron 7 splicing in the CBE-edited *sot5* background plants (A) and CBE-edited WT background plants (B). IR indicates the transcript retained intron 7. -22 ss indicates the transcript derived from -22 cryptic splice site. +1ss indicates the transcript derived from the authentic splice site in WT. (C) qPCR analysis of splicing efficiency of *SOT5* intron 7 in *E61*, *sot5 EX8-CR-2,* and *sot5-IN7-CR* plants. The data in *sot5* were set as 1. (D) qPCR analysis of splicing efficiency of *SOT5* intron 7 in WT *IN7-CR* and WT *EX8-CR* plants. The data in WT were set as 1. The data are means of three biological replicates (bars indicate SD). Asterisks indicate significant differences between mutant and control (Student’s *t* test). **, *P* < 0.01.

To determine whether the exon 8 ESE is generally required or only functions in the context of a compromised splice site, we used CBE to introduce the same synonymous G-to-A mutation in the WT background and generated homozygous WT *EX8-CR* edited plants. In contrast to the *sot5* mutant, the mutation of this ESE element in WT plants (WT *EX8-CR*) had no effects on leaf color and plant growth (Figure 3B and 3C), and did not cause a significant reduction in the splicing efficiency of intron 7 (Figure 4B and 4D), indicating that the ESE functions specifically in the *sot5* background. Interestingly, a G-to-A mutation at the 3′ splice site (3′ss) of *SOT5*intron 7 in WT background (WT *IN7-CR*), led to a dramatic reduction in the mature +1ss transcript, as detected by specific qPCR primers (Figure 4D). Yet semi-quantitative RT-PCR still detected a similar mature transcript (Figure 4B). Sequencing of the PCR product of this similar mature transcript in WT *IN7-CR* revealed a 9-nucleotide deletion at the 5′ end of exon 8—a feature distinct from the mature transcript in WT.

Taken together, these results indicate that the AAGAAGA motif in exon 8 functions as a bona fide ESE, which is essential for the efficient splicing of intron 7 only in *sot5*, but not in WT. In addition, the *SOT5* transcript retaining intron 7 encodes a functional SOT5 protein that can functionally substitute for native SOT5 protein in vivo under normal growth conditions.

### The AAG repeat motif is essential for the ESE function in efficient intron 7 splicing

To finely evaluate the role of the AAGAAGA sequence in exon 8 ESE, we performed a detailed mutational analysis and generated a series of *SOT5 IN7* CDS constructs driven by the 35S promoter, each containing specific nucleotide substitutions within the ESE element (Figure 5A). These constructs were transformed into the *sot5* mutant to assess their impact on intron 7 splicing efficiency. Our results showed that all transgenic lines exhibited a WT-like phenotype and rescued the virescence of the *sot5* mutant under normal growth conditions (Figure 5B). qPCR analysis revealed that, compared to the control M1 (*SOT5 IN7* CDS containing AAGAAGA), all single-base mutations, such as M2 (AAGAAAA), M3 (AAAAAGA), M4 (AAAAAAA) and M5 (AAGAACA), or multiple-base mutation, such as M6 (CTCTCTC) caused a significant reduction in intron 7 splicing efficiency (Figure 5A, 5C and 5D). These findings demonstrate that the conserved ESE sequence is essential for promoting efficient intron splicing.

**Figure 5.**
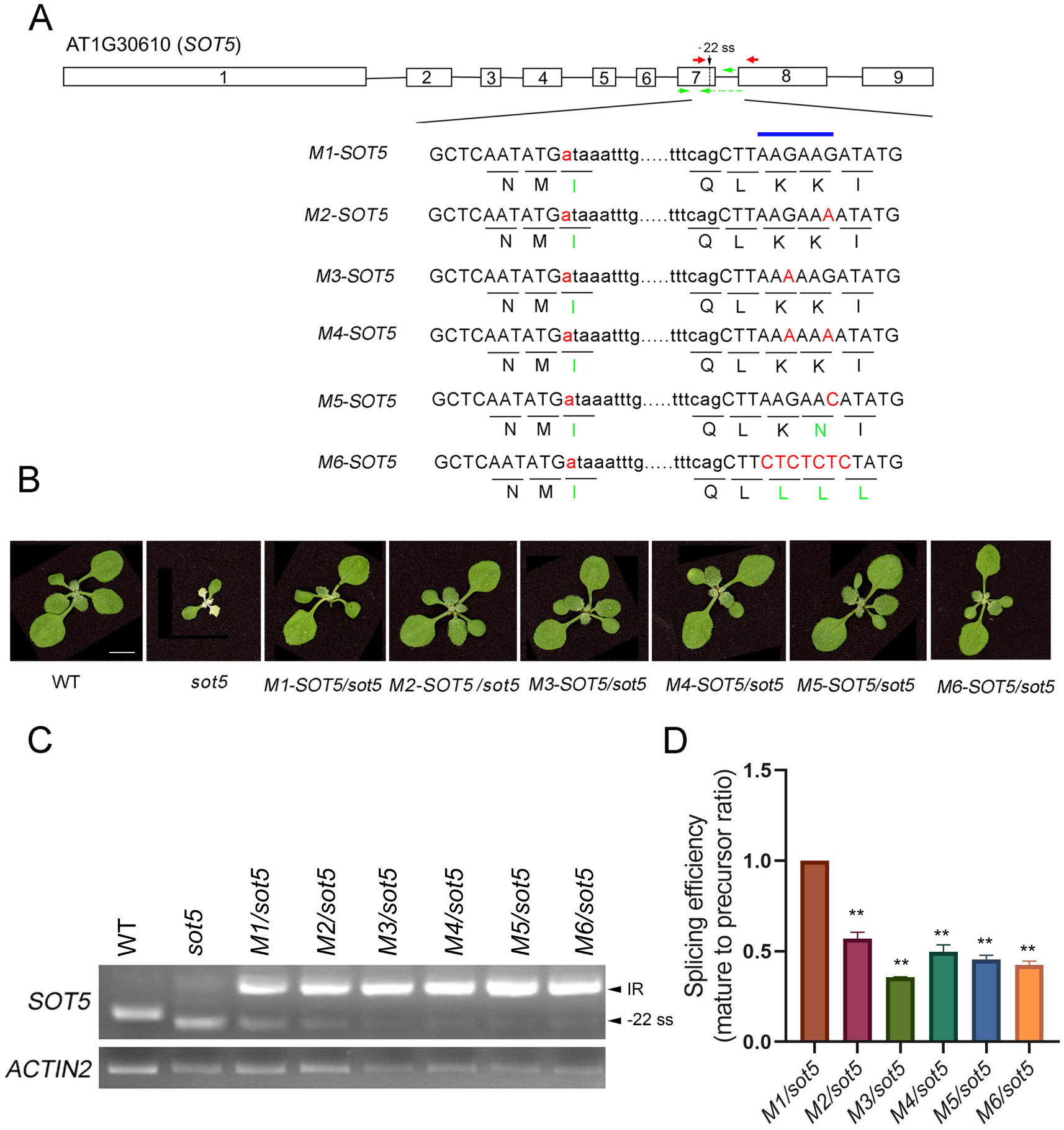
The ESE motif is essential for efficient intron7 splicing. (A) *SOT5* gene structure with targeted mutations in ESE. Red, mutation sites; blue bar, ESE location. Corresponding amino acid sequences are shown below, with green indicating altered residues. Red and green arrows indicate primers used for RT-PCR and RT-qPCR analysis, respectively. (B) Phenotypes of T1 transgenic plants expressing mutated ESE constructs depicted in (A). WT and *sot5* are included as controls. Bar = 1 cm. (C) Semi-quantitative PCR analysis of *SOT5* intron7 splicing in the transgenic plants. (D) qPCR analysis of splicing efficiency of *SOT5* intron 7, using the *M1/sot5* plant as the reference (set as 1). Data represent means ± SD of three biological replicates. Asterisks indicate significant differences between mutant and control (Student’s *t* test). **, *P* < 0.01.

### Mutations in *SR45* rescue the *sot5* phenotype by reducing intron 7 splicing efficiency

In animals, ESEs enhance splicing of their nearby weak splice sites through binding to one or more SR or SR-like proteins, which in turn recruit U1 snRNP and U2 snRNP to initiate the spliceosome assembly. We therefore hypothesized that the ESE element in exon 8 binds one or more SR or SR-like proteins, and this interaction is disrupted in *E61*, leading to impaired splicing. Based on known RNA-binding specificities, we prioritized several candidates likely to recognize the AAGAAGA motifs, including AtSR45 (Xing et al., 2015; Fanara et al., 2025) and its homologs SR45a and another canonical SR protein, SR30 (Tacke and Manley, 1995). One effective approach is to generate double mutants of *sr* and *sot5*. If a loss of a particular SR protein rescues the *sot5* phenotype, which is similar to the effect of the *E61* mutation, it would indicate that this protein probably binds to the AAGAAGA motif.

To know the roles of these factors in regulating intron splicing efficiency, we first identified two T-DNA insertion mutants, *sr45a* and *sr30*, from the Arabidopsis Biological Resource Center (Figure S2), and then generated their double mutants with *sot5* via crossing. Unfortunately, neither *sr45a* nor *sr30* rescued the *sot5* phenotype (Figure 6A and Figure S2C), suggesting that SR45a and SR30 is not involved in recognition of the exon 8 ESE. Then, we used CRISPR/Cas9 technique to knockout *SR45* and *SR45a* in the *sot5* background. Interestingly, the *sr45 sot5* double and *sr45 sr45a sot5* triple mutants displayed a robust rescue of the virescent leaf phenotype, whereas the *sr45a sot5* double mutant resembled *sot5* (Figure 6A and 6B), indicating that mutations of *SR45* alone is sufficient to rescue the *sot* phenotype. Consistently, the chlorophyll content and *F*_v_/*F*_m_ values of *sr45 sot5* were comparable to those of WT and significantly higher than those of *sot5* (Figure 6C-E). In addition, PCR analysis showed a significantly reduced splicing efficiency of *SOT5* intron 7 in both the *sr45 sot5* double and *sr45 sr45a sot5* triple mutants, similar to those in *E61* (Figure 6F, 6G). Thus, our data suggest that SR45 is a *trans*-acting factor essential for the function of the *SOT5* exon 8 ESE by binding to this ESE specifically.

**Figure 6.**
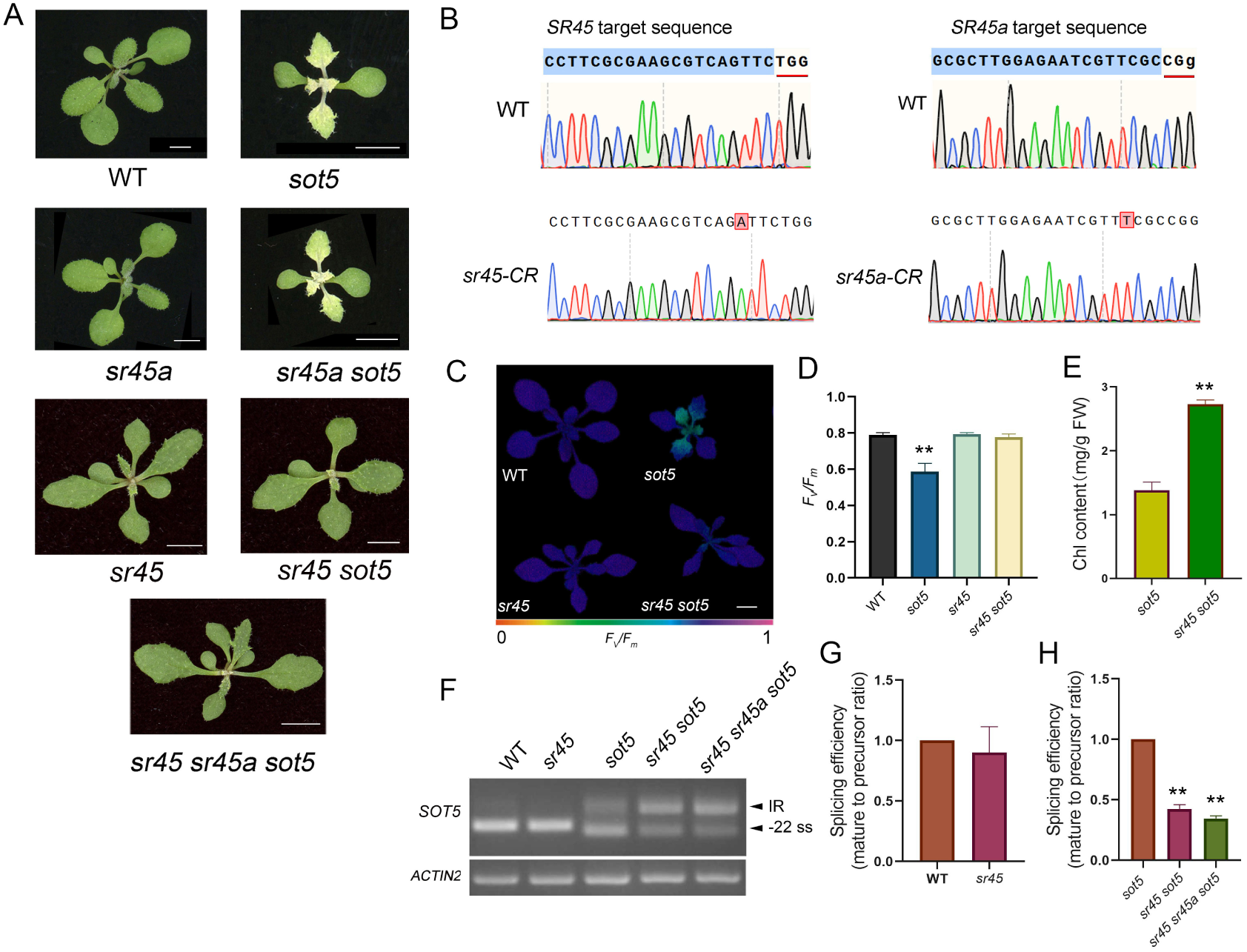
*SR45* mutations rescue the *sot5* phenotype via reducing intron 7 splicing efficiency. (A) Phenotypes of 20-day-old WT and single, double and triple mutant plants. (B) Sanger sequencing of the target regions confirmed the specific genomic edits in the *sr45-CR* and *sr45a-CR* plants. The red lines highlight the PAM sequences adjacent to the edited sites. (C) *F*_v_/*F*_m_ images of *sr45 sot5* compared with WT, *sr45* and *sot5.* (D, E) Chlorophyll content and *F*_v_/*F*_m_ values of *sr45 sot5* compared with WT, *sr45* and *sot5.* (F) Semi-quantitative PCR analysis of *SOT5* intron7 splicing in *sr45 sot5* and *sr45 sr45a sot5* mutants. (G) qPCR analysis of *SOT5* intron 7 splicing efficiency in WT and *sr45*. The splicing efficiency in *sot5* line was set to 1 as reference. (H) PCR analysis of *SOT5* intron 7 splicing efficiency in *sot5*, *sr45 sot5* and *sr45 sr45a sot5* mutants. The splicing efficiency in *sot5* was set to 1 as reference. Data represent means ± SD of three biological replicates. Asterisks indicate significant differences between mutant and control (Student’s *t* test). **, *P* < 0.01.

### Mutating the proximal ESEs of *GPDHC1* intron2 mimicked its splicing defects caused by SR45 mutation

To identify additional ESE regulated by SR45, we did the literature mining regarding to the transcriptome data of *sr45* mutant (Xing et al, 2015; Zhang et al, 2017). And we also performed RNA-seq on *sot5* and *sr45 sot5* mutants. Transcriptomic analysis (Table S2) combined with RT-PCR verification revealed that several introns were significantly retained in *sr45 sot5*, compared to *sot5* (Table S3 and Figures S4). The reported SR45 RIP data demonstrated that these mis-spliced RNAs in the *sr45* mutant are also bound by SR45-GFP (Xing et al., 2015), suggesting they are likely direct targets of SR45 (Table S3). Analysis of the exon sequences flanking these retained introns revealed a frequent occurrence of AAG-like motifs within 50 nucleotides upstream or downstream of the splice sites (Table S3). We then selected intron 2 of *GPDHC1* for functional validation, as it is alternatively spliced in WT and *sot5* but largely retained in the *sr45* mutant background (Figure 7A and 7B). This intron is bordered by the AAG repeat motifs in exon 2 and exon 3 (Figure 7C). To examine the function of these motifs, we transformed both native and mutated versions of the AGG-motif within the genomic *GPDHC1* driven by the 35S promoter into WT plants. Analysis of T_1_ plants showed that the native *GPDHC1* transgene (*N-GPDHC1/WT*) recapitulated the alternative splicing pattern, whereas the AAG-mutated version (*M-GPDHC1/WT*) resulted in dramatic decrease of intron 2 splicing (Figures 7D, 7E). These results indicate that SR45-dependent splicing of *GDPHC1* intron 2 is mediated by the ESE elements. Notably, as intron 2 resides within the 5′UTR, its retention leads to an elongated 5′UTR that could subsequently affect protein translation efficiency or mRNA stability. It is interesting that *GPDHC1* encodes Cytosolic Glycerol-3-Phosphate Dehydrogenase 1 that is critical for maintaining cellular NADH/NAD+ balance. It is a central enzyme in managing cellular energy metabolism and redox balance, especially under stressful environmental conditions. The Arabidopsis *gpdhc1* mutant exhibits sensitivity to salt stress and ABA (Shen et al., 2006), which aligns with some phenotypes observed in the *sr45* mutant, indicating that *GPDHC1* is an important target of SR45.

**Figure 7.**
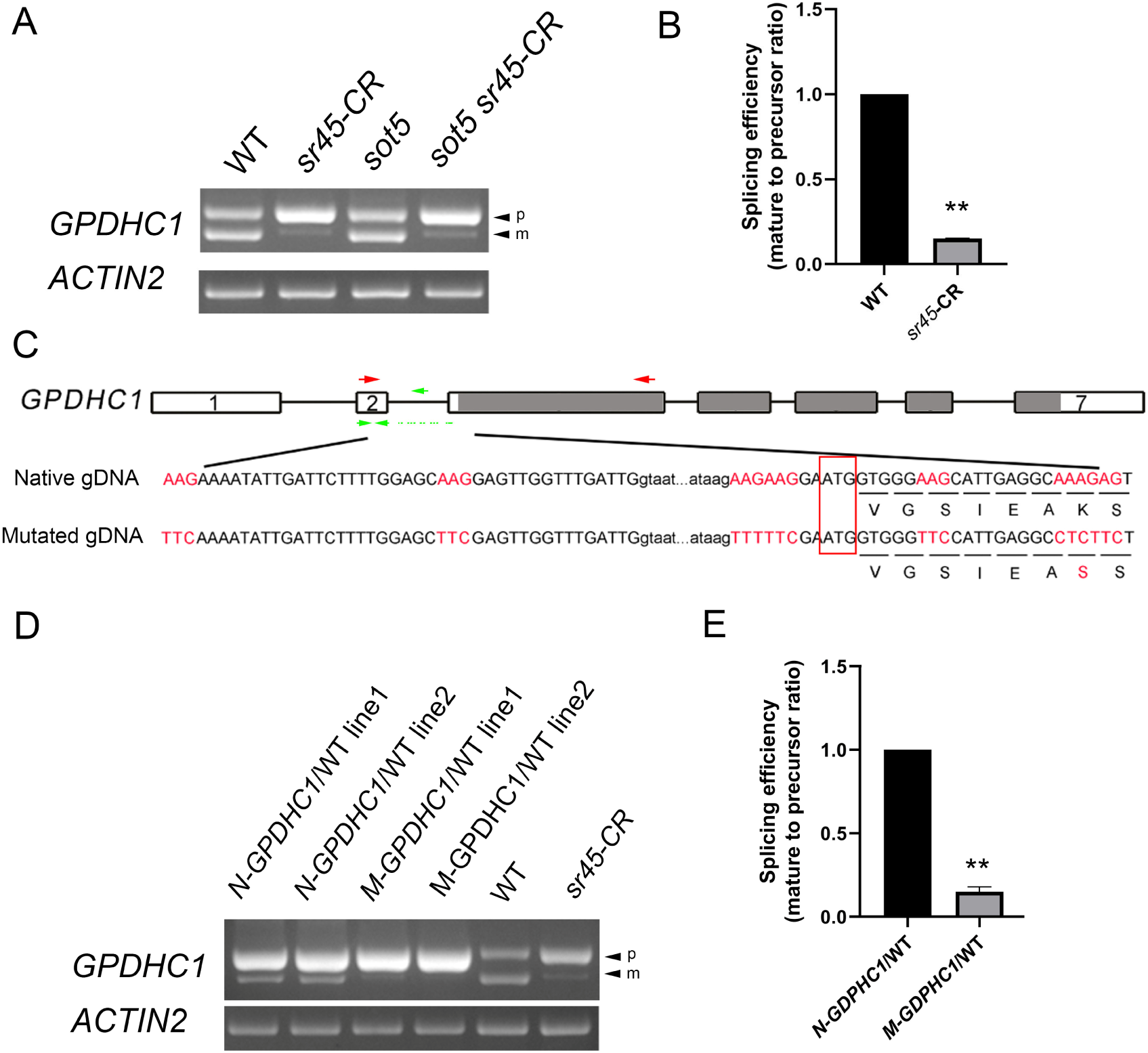
Mutating the proximal ESEs of *GPDHC1* intron2 mimicked its splicing defects caused by SR45 mutation. (A) PCR analysis of *GPDHC1* intron 2 splicing in WT, *sot5*, *sr45-CR*, and *sot5 sr45-CR* plants. (B) Splicing efficiency of *GPDHC1* intron 2 was analyzed by RT-qPCR. Data represent means ± SD of three biological replicates. The splicing efficiency in WT was set to 1 as reference. (C) Schematic diagram of *GPDHC1* gene structure with AAG repeat sequences highlighted in red. These putative exonic splicing enhancer (ESE) motifs are located on exons adjacent to intron 2, all falling within 50 bp from its splice junctions. White rectangles indicate 5’ and 3’UTRs, gray rectangles indicate coding regions, and black lines indicate introns. Red and green arrows indicate primers used for RT-PCR and RT-qPCR analysis, respectively. (D) RT-PCR analysis of T1 transgenic plants expressing the native *GPDHC1* transgene (*N-GPDHC1*) and the AAG-mutated version (*M-GPDHC1*). Intron 2 splicing patterns in WT and *sr45-CR* was used as controls. (E) Splicing efficiency of *GPDHC1* intron 2 in (D) was quantified by Image J software. The splicing efficiency in *N-GPDHC1* transgenic line was set to 1 as reference. Asterisks indicate significant differences between mutant and control (Student’s *t* test). **, *P* < 0.01.

### AtSR45 exhibits stronger affinity to RNA containing the AAG repeat motif, rather than AAA repeat motif, in vitro

To determine whether SR45 can directly bind to the AAGAAG motif, we performed RNA electrophoretic mobility shift assays (REMSA) with the purified N-terminal MBP-tagged SR45 protein from bacteria and the Cy5-labeled RNA probe containing three tandem AAGAAG repeats (6×AAG). REMSA results showed that MBP-SR45 directly bound the Cy5-labeled 6×AAG RNA probe (Figure 8A). This binding was effectively competed by an unlabeled “cold” probe but not by a mutant probe containing six AAA repeats (6×AAA) (Figure 8B). These results revealed that SR45 binds directly to RNA containing the AAG repeat motif, and exhibits significantly weaker binding affinity toward the AAA repeat motif RNA. Our finding is also consistent with the following reported data: SR45 binds the AAGAAG RNA motif in both intron-containing and intronless genes revealed by RIP-seq experiments (Xing et al., 2015). RRM domain of SR45 specifically recognizes AGAAG motifs via three key residues (H101, H141, and Y143) demonstrated by SELEX assays (Fanara et al., 2024).

**Figure 8.**
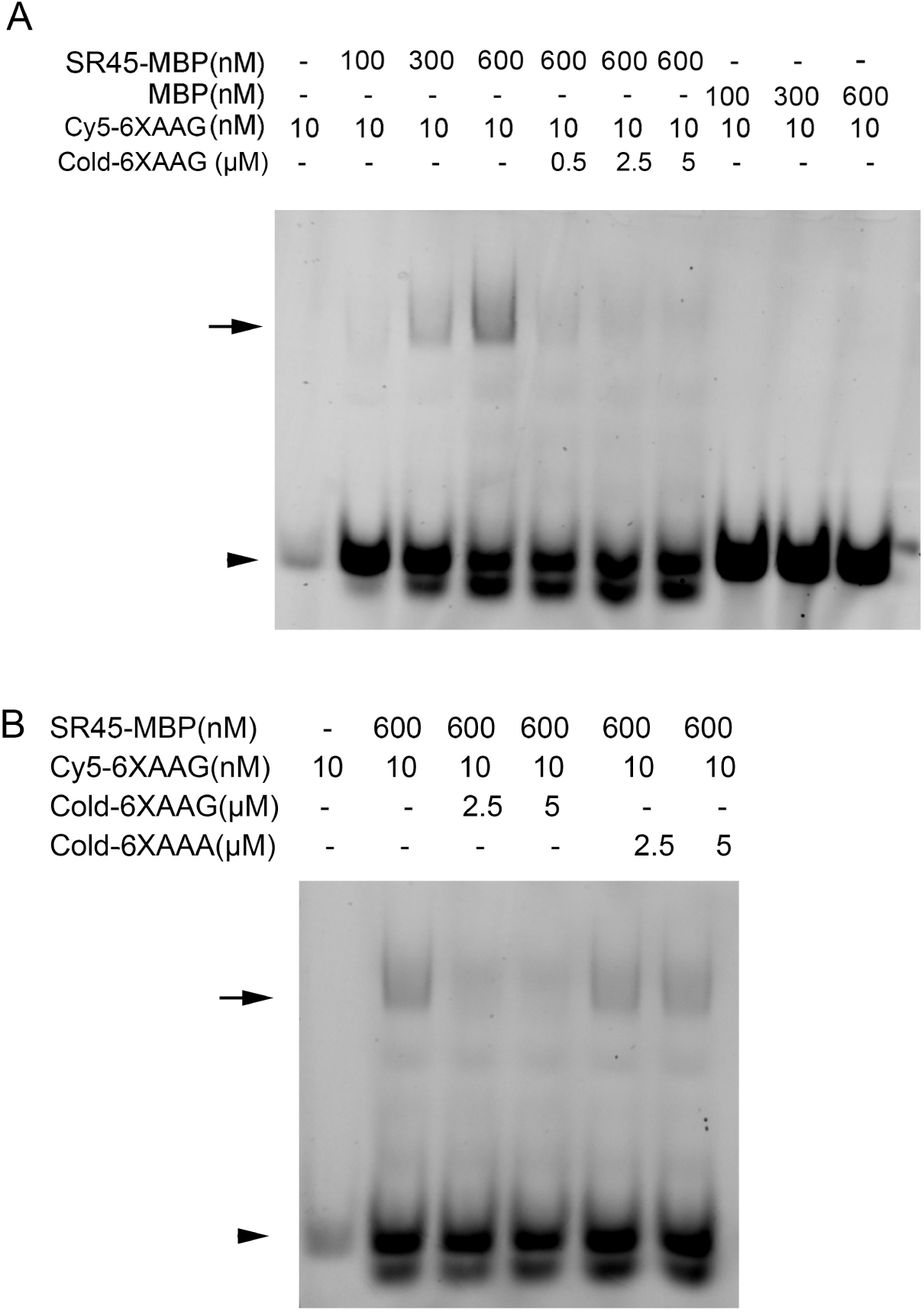
SR45 specifically binds to the RNA containing AAG repetitive motif. (A) REMSA assays of interaction between MBP-SR45 and the Cy5-labeled 6×AAG RNA probe. MBP protein alone was used as a negative control. Cold-6×AAG means the unlabeled probe competing with the labeled probe. (B) Competitive REMSA analysis of MBP-SR45 binding to the Cy5-labeled 6×AAG RNA in the presence of unlabeled 6×AAG RNA or a mutated probe containing six AAA repeats (cold-6×AAA).

## Discussion

### A model for SR45-dependent splicing regulation via binding ESE motifs

In this study, our screening for and characterization of *sot5* suppressors reveal that AAG repeat sequences downstream of an alternative splice site function as ESEs bound by SR45. First, intragenic mutations causing a significant decrease in the splicing efficiency of intron 7 suppress *sot5* leaf virescence; second, the intragenic mutations lying within an ESE change splicing patterns of an intron with weak splice sites but not with constitutive splice sites; third, loss-of-function mutations in *SR45* rescue the *sot5* phenotype, mimicking the ESE mutation; last, SR45 specifically binds to the ESE motif to promote efficient intron splicing. Based on these findings, we propose a working model for how SR45 regulates splicing efficiency through binding to ESE (Figure. 9). In the WT background, the canonical 5′ss of *SOT5* intron 7 is strong and efficiently recognized by U1 snRNP, resulting in constitutive splicing. Although an ESE motif exists nearby in exon 8, it exhibits little or no ESE activity, as its mutation does not have an impact on intron 7 splicing (Figure 9A). In the *sot5* mutant, a GT-to-AT mutation at this canonical 5′ss disrupts U1 snRNP binding. This inactivation leads to two outcomes: approximately 40% of transcripts retain intron 7, while the remaining transcripts activate two cryptic splice sites, a dominant site at -22 nt (∼50%) and a weaker site at +10 nt (∼10%). As we previously showed that the -22 nt site is a weak splice site and requires assistance from U1 snRNP-associated proteins RBP45d and PRP39a (Figure 9B). This study demonstrates that efficient usage of at the -22 nt cryptic site also strictly depends on SR45. Loss of SR45 or mutation of its binding ESE results in a dramatic shift toward intron 7 retention (Figure 9C). Given that SR45 is known to interact with U1-70K and U2AF65 (Day et al., 2012), we propose a mechanism by which SR45 binds to the ESE proximal to intron 7 and facilitates intron excision by simultaneously bridging interactions with both U1 and U2 snRNPs.

**Figure 9.**
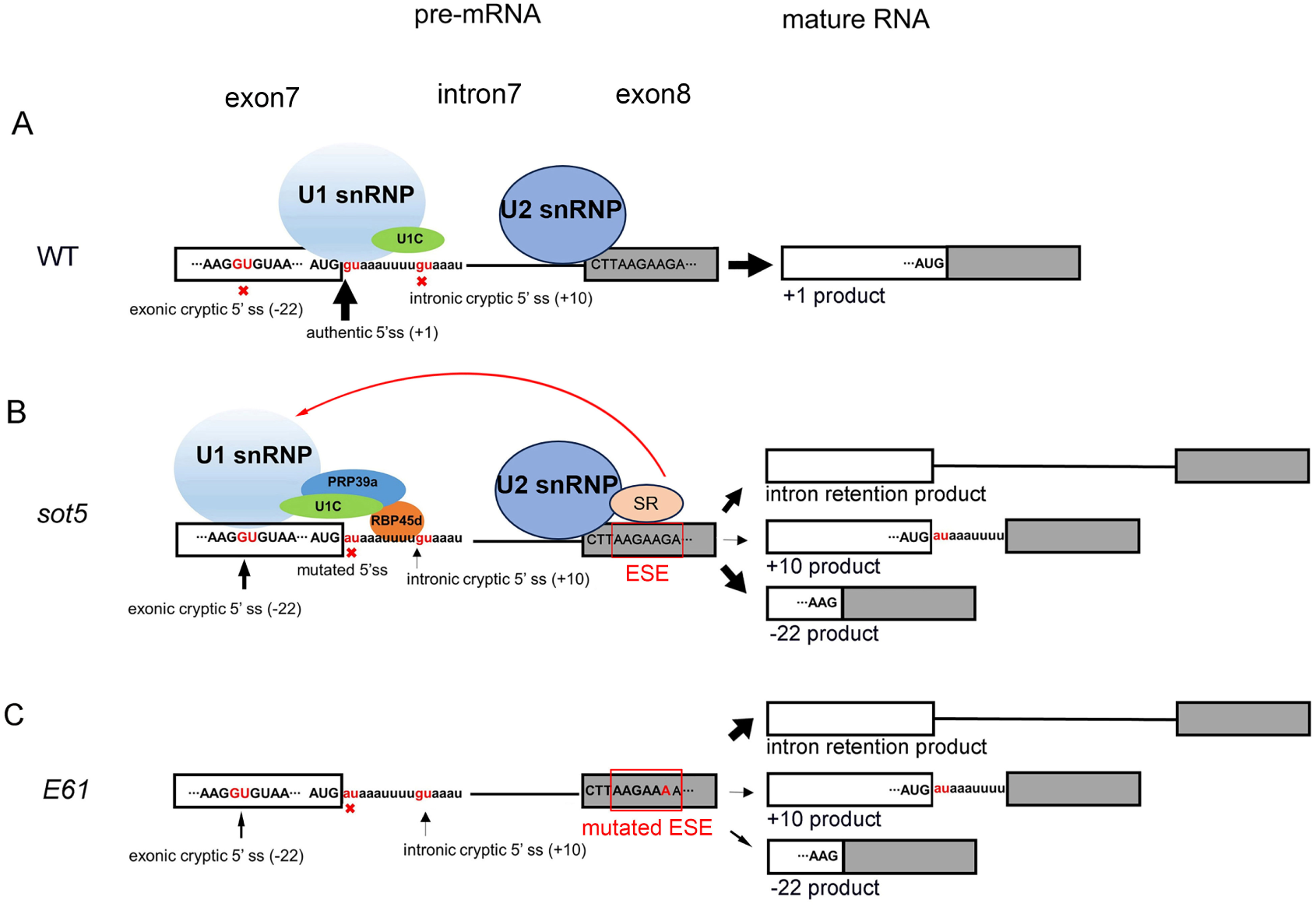
Working model of SR45-mediated enhancement of intron splicing via ESE binding. (A) In WT plant, the strong canonical 5′ss of *SOT5* intron 7 is efficiently recognized by U1 snRNP, resulting in constitutive splicing. Although an AAG repeat sequence exists nearby in exon 8, it exhibits little or no ESE activity. (B) In the *sot5* mutant, a GT-to-AT mutation at the canonical 5′ss disrupts its recognition by U1 snRNP, leading to two outcomes: approximately 40% of transcripts retain intron 7, while the remaining transcripts activate two cryptic splice sites: the -22 nt site (∼50%) and the +10 nt site (∼10%). (C) This study demonstrates that splicing at -22 nt site requires additional splicing factor SR45. Mutations in SR45 and its cognate ESE results in increased intron 7 retention. Based on known interactions between SR45 and U1-70K (U1 snRNP component) and U2AF35 (U2 snRNP component), we propose that SR45 binds to the ESE proximal to intron 7 and facilitates intron splicing by simultaneously bridging interactions with both U1 and U2 snRNPs.

### A probable pathway for the evolution of alternative splicing

We found that AAG repeats proximal to strong, constitutive splice sites are functionally inert, whereas the same sequences adjacent to weak, cryptic splice sites can be co-opted as functional ESEs. The *sot5* suppressor screen illustrates a potential evolutionary route for AS events. Generally, a strong, constitutive splice site mask weak, cryptic site. Once a constitutively spliced intron mutates, cryptic splice sites are formed with proximal intronic or exonic sequences evolving into *cis*-regulatory elements such as ESEs, ESSs, ISEs, or ISSs. These elements are recognized by specific *trans*-acting factors including SR and hnRNP proteins to promote efficient splicing at these weak sites, thereby converting constitutive splicing (CS) into AS.

This regulatory landscape of AS involves not only the conserved core splicing signals and factors but also additional *trans*-acting factors, making it more complex and competitive between weak splice sites. Each weak site is often associated with a unique sequence motif and a specific binding factor. For example, a U-rich intronic splicing enhancer (ISE) downstream of the –22 nt site is bound by RBP45d, which promotes usage of this site over the +10 nt site. It is possible that RBP45d binding to the U-rich sequence within the +10 nt site could actively block its recognition, and mutations of RBP45d de-represses the +10nt site. We observed that either mutations in SR45 or its associated ESE block the recruitment of U1 and U2 snRNPs to nearby sites, resulting in inefficient usage of both cryptic splice sites. The interplay among alternative splicing factors can be cooperative or antagonistic. For instance, SR45 and RBP45d coordinately promote usage of -22 nt site usage. Conversely, the disruption of an ESE may not merely abolish enhancement but could potentially create a silencer (ESS) that recruits repressive hnRNP proteins. This highlights the intricate and context-dependent interplay among splicing factors.

### Validation of ESE function relies on experimental approaches

The regulation of AS sites is highly complex and context-dependent. A single splice site within a gene is often regulated by multiple SR and hnRNP proteins, which can act cooperatively or antagonistically (Chen and Manley, 2009). Because the sequences surrounding each alternative splice site are unique, and RNA-binding proteins exhibit specific sequence affinity, predicting functional elements computationally remains challenging. Therefore, functional validation through experimentation is indispensable. This is exemplified in human therapeutics, where the development of small-molecule drugs for human diseases such as SMA relies on detailed studies of alternative splicing regulation in the *SMN2* gene (Hofmann et al., 2000; Wan and Dreyfuss, 2017; Campagne et al., 2019). This process involves large-scale experimental screening for oligonucleotides that enhance *SMN2* splicing, ultimately leading to optimized antisense oligonucleotides (ASOs) that are developed into therapeutics (Wan and Dreyfuss, 2017). Notably, ASO drugs designed for different mutation sites in *SMN2* or *DMD* are not universally applicable (Wan and Dreyfuss, 2017; McNally and Wyatt, 2017). To identify alternative splice sites and their regulatory elements, highly relevant to human splicing-related diseases, computational methods have been developed to predict splice sites and ESEs in human (Blencowe, 2000; Fairbrother et al., 2002; Cartegni et al., 2003; Wang et al., 2004; Jaganathan et al., 2019; Dawes et al., 2022). These methods are essentially empirical, relying on compiled data from disease-associated mutations and human RNA-seq databases. Many studies experimentally validate splice sites and regulatory elements using mini-gene constructs carrying mutated splicing elements.

Although prediction of ESEs has been reported in Arabidopsis (Pertea et al., 2007), functional validation is scarce. Direct cloning of plant genes into binary vectors for stable transformation provides an effective system for studying splicing elements, capturing regulatory features that may be absent in mini-genes. For crop species, where genetic transformation is less efficient, developing reliable transient transfection systems for mini-gene splicing reporters is critical prerequisite for functional studies. Once validated, genome editing techniques can be applied to modify specific sequences for crop breeding, significantly reducing time and resource costs.

### SR45-regulated ESEs are potential targets for precision crop improvement

In this study, we extended our validation to intron 2 of *GPDHC1*, another SR45 target, confirming that mutation of its proximal ESE similarly impairs splicing. This underscores the general utility of AAG repeat motifs as SR45-responsive ESEs and reveals a broader mechanism for post-transcriptional regulation.

Our findings provide a promising strategy for precision crop breeding through gene editing. Fine-tuning gene expression is often critical for optimizing agronomic traits. For example, *Waxy* (*Granule-Bound Starch Synthase 1*, *GBSS1*) expression level is significantly associated with cooking and eating quality in rice. Beyond modulating promoters or coding sequences (Huang et al., 2021; Xu et al., 2021; Zhang et al., 2022), regulating mRNA splicing efficiency via *cis*-regulatory elements offers another powerful strategy. Editing pleiotropic *trans*-acting factors such as SR proteins is often problematic due to unintended effects on numerous traits. In contrast, precisely modifying *cis*-regulatory elements within target genes allows for fine-tuning its expression without pleiotropy. This approach has been applied in human gene therapy for DMD (Qiu et al., 2023), demonstrating the broad prospects for the treatment of genetic diseases. We propose a general workflow for crop improvement through editing splicing regulatory elements. First, determining whether genes controlling key agronomic traits undergo alternative splicing. Second, identifying the *trans*-acting factors regulating these splicing events through literature mining or experiments. Third, mapping the corresponding cis-regulatory elements (ESE, ESS, ISE, ISS etc.). Fourth, using transient gene expression system to test the effect of mutations on splicing efficiency. Last, introducing precision mutations in elite varieties and evaluating their effects on splicing and agronomic performance. This pathway from mechanistic discovery to validated editing strategy can significantly reduce the time and resource costs associated with developing improved crop varieties.

## Supplemental Data

**Figure S1.**
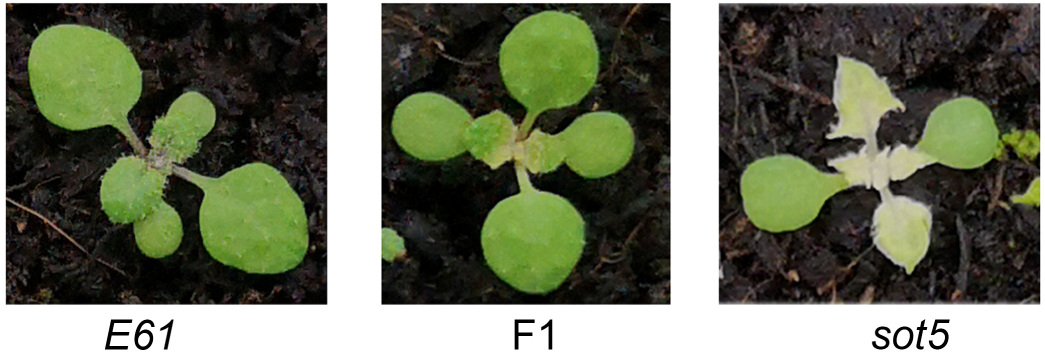
Phenotype of the F1 progeny of the *E61* × *sot5* cross.

**Figure S2.**
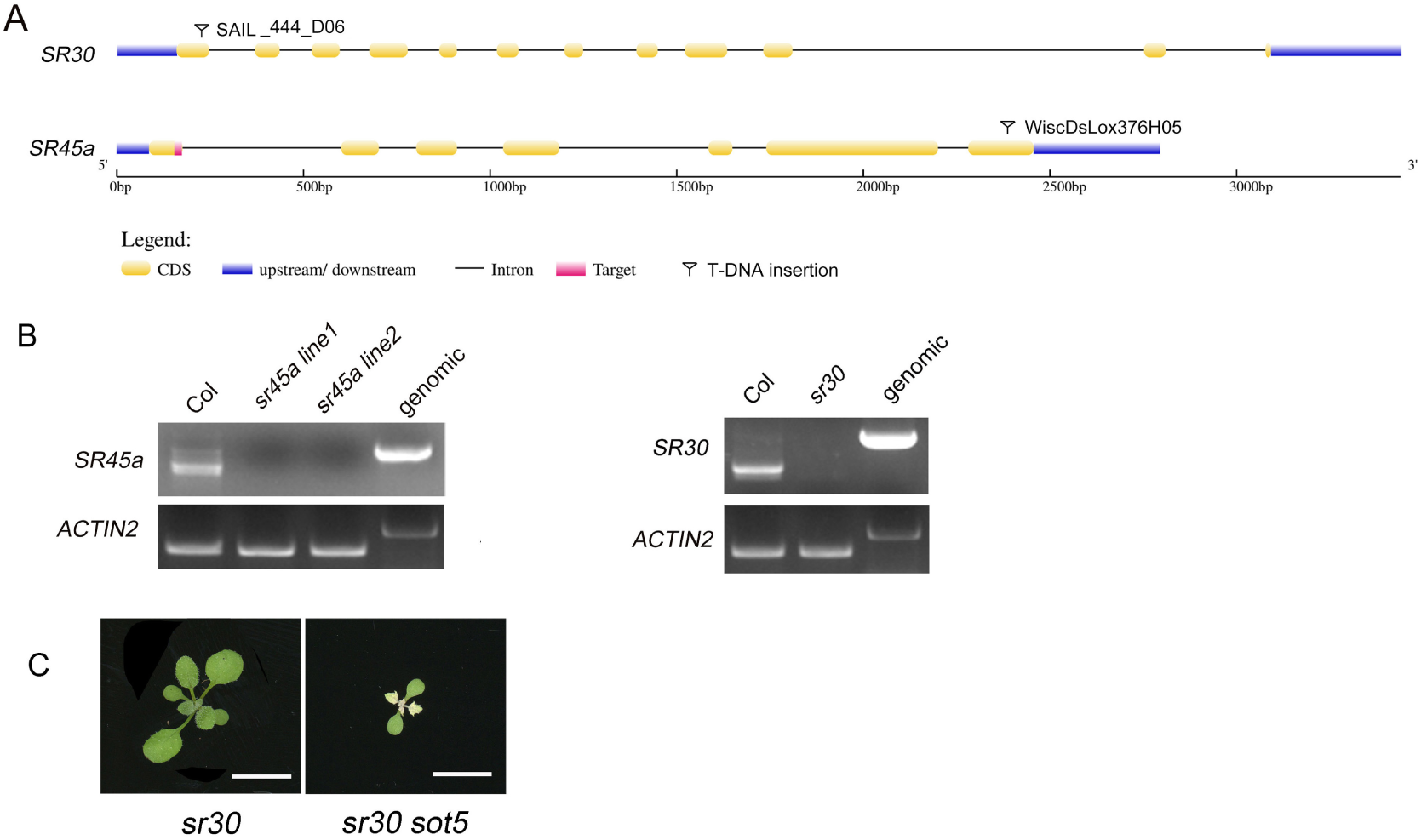
Identification of *sr45a* and *sr30* T-DNA insertion mutants and phenotype of *sr30 sot5* double mutant. (A) The gene structure of *SR45a* and *SR30* with the positions of the T-DNA insertion. (B) RT-PCR analysis confirmed the absence of the transcripts of *SR45a* and *SR30* in the mutants. (C) Phenotype of the *sr30* and *sr30 sot5* double mutant.

**Figure S3.**
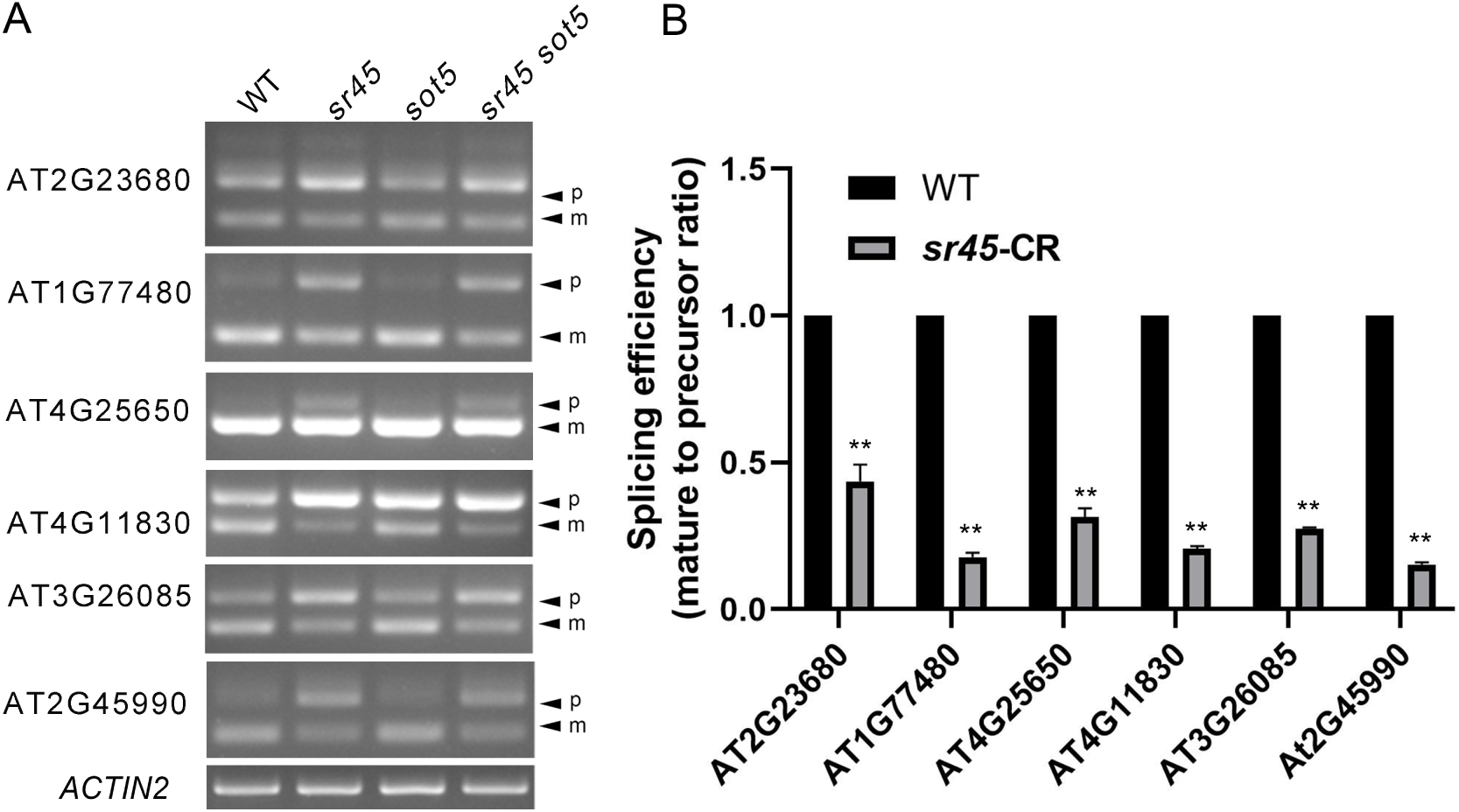
Validation of the introns retained in the *sr45* background by RT-PCR and RT-qPCR. (A) RT-PCR analysis of the selected introns (listed in Table S3) which are significantly retained in *sr45* background and their transcripts are simultaneously associated by SR45-GFP. (B) Splicing efficiency of the above introns was analyzed by RT-qPCR. Data represent means ± SD of three biological replicates. Asterisks indicate significant differences between mutant and control (Student’s *t* test). **, *P* < 0.01.

## Supplemental Tables

**Table S1.** Segregation analysis of the F2 population from the *E61 × sot5* cross.

**Table S2.** Introns retained in *sr45 sot5* mutant.

**Table S3.** The exon sequences flanking the retained introns in *sr45* background contain the AAG-like sequences.

**Table S4.** Primers used in this study.

**Table S5.** Gene accession numbers in this study.

## Material and Methods

### Plant Materials and Growth Conditions

The Arabidopsis (*Arabidopsis thaliana*) ecotype Columbia-0 (Col-0) was used as the wild type (WT) in this study. The mutants *sot5* were previously described (Huang et al., 2018). The *sr45a* (WiscDsLox376H05) and *sr30* (sail_444_D06) mutant was obtained from the ABRC stock center (https://abrc.osu.edu/). Double mutants *sr45a sot5* and *sr30 sot5* were identified from F_2_ generations derived from crosses between single mutants by the PCR-based genotyping procedure. The primers used for genotyping are listed in Table S5. *sr45 sr45a sot5* triple mutant was obtained by editing *sr45* and *sr45a* in *sot5* background using CRSIPR/Cas9 techniques described previously (Huang et al, 2022). *sr45 sot5* double mutant was obtained by editing *sr45* in *sot5* background. Seeds were surface-sterilized by 75% ethanol and stratified at 4°C for 3 days, and then sown onto half-strength Murashige and Skoog (MS) medium with 1% sucrose. Seeds were also directly sown in soil and grown in a phytotron with long-day conditions (16 h light/8 h dark) and light intensity (100 μmol photons m^-2^ s^-1^) at 22°C.

### Plasmid Construction and Transformation

*E61* gDNA, *E61* CDS, and *SOT5 IN7* CDS sequences were cloned into the pENTR SD/D-TOPO entry vector, and then recombined into the pGWB2 destination vector. *M1∼M5-SOT5* CDS with different mutations were cloned into pCambia1300-35S-NOS vector. The above vectors were transformed into *sot5* mutants. The native and mutated *GPDHC1* gDNA sequence were cloned into the pENTR SD/D-TOPO entry vector, and then recombined into the pGWB5 destination vector and were transformed into Col-0.

### Generation of WT *IN7-CR, sot5 IN7-CR,* WT *EX8-CR* and *sot5 EX8-CR* mutant lines by CBE

We constructed the Cytosine Base Editors, AtCBE-NGG and AtCBE-NG for dicot plants by replacing the OsU6 and ZmUBI promoters in the rice editors evoFERNY and evoFERNY-NG (Wei et al., 2022) with the AtU6 and RPS5A promoters, respectively. The resulted AtCBE-NGG-RPS5A-Ferny vector was employed to target the ESE motif in *SOT5* exon 8, while the resulted AtCBE-NG-RPS5A-Ferny vector was used to edit the 3’ splice site (3’ss) of *SOT5* intron 7. sgRNAs were designed based on the CBE vector specifications and the target sites. These constructs were subsequently transformed into either WT or *sot5* mutant backgrounds. Homozygously edited plants were identified in T_1_ and T_2_ generations through PCR genotyping and sequencing.

### RNA extraction, RT-PCR, RT-qPCR and sequencing of the RT-PCR products

Total RNAs were extracted from seedlings or different plant tissues by RNA Easy Fast Plant Tissue Kit (TIANGEN, DP452) according to the manufacturer’s instructions. Reverse transcription was conducted by PrimeScript RT reagent Kit with gDNA Eraser (Takara, RR047A). Semi-quantitative RT-PCR was carried out using the gene-specific primers (Table S1). qPCR was previously described (Zhang et al., 2021). To identify the splicing variants, the PCR products amplified by the primers spanning the introns were purified for sanger sequencing.

### RNA-seq and Bio-informatic analysis

rRNA-depleted RNA-seq experiments and subsequent analysis was performed on *sot5* and *sr45 sot5* mutant plants by Shanghai OE Biotech. Differential splicing analysis was performed using rMATS (Shen et al., 2014).

### Protein expression, purification and RNA electrophoretic mobility shift assay (REMSA)

The AtSR45 CDS was cloned into the expression vector pMal-c5x, and protein expression was induced using 0.5 mM isopropyl-b–D–thiogalactopyranoside at 16°C for 12 h in *E. coli* strain Rosetta. The protein was purified by NEB Amylose Resin according to the manual (NEB, E8021S). The RNA probes were chemically synthesized, and their 5’-end were labeled by Cy5 (Sangon Biotech (Shanghai) Co., Ltd). The sequences of the probes were as follows:

Cy5-6XAAG: 5’-(CY5) AAGAAGAAGAAGAAGAAG-3’;

The corresponding non-labeled and mutated probes were also chemically synthesized for competition assays.

Cold-6XAAG: 5’- AAGAAGAAGAAGAAGAAG -3’

Cold-6XAAA: 5’- AAAAAAAAAAAAAAAAAA -3’

For REMSA assay, AtSR45 or AtSR45a protein and RNAs were incubated at room temperature for 30 min in the reaction solution (10 mM HEPES pH 7.3, 20 mM KCl, 1 mM MgCl2, 1 mM dithiothreitol, 5% glycerol (v/v) and 0.1 μg tRNA). The reactants were analyzed using a native polyacrylamide gel, and signals were detected by Azure Biosystems C600.

### Accession Numbers

The accession numbers of the gene described in this study are listed in Table S5.

## ACKNOWLEDGMENTS

We thank the ABRC for providing the *sr45a* (WiscDsLox376H05) and *sr30* (sail_444_D06) seeds. This work was supported by the National Natural Science Foundation of China (32171293), The Fund of Innovation Program of Shanghai Municipal Education Commission (2021-01-07-00-02-E00117), and Shanghai Engineering Research Center of Plant Germplasm Resources (17DZ2252700).

## AUTHOR CONTRIBUTIONS

W.H. conceived and designed the research; Y.Z W.S., L.Z., F.L. K.Y. J.W. H.Z. and W.H. performed the experiments and analyzed the data; W.H. and J.H. supervised the experiments and wrote the article.

## Notes

### Competing Interest Statement

The authors have declared no competing interest.

